# Floating frogs sound larger: environmental constraints on signal production drives call frequency changes

**DOI:** 10.1101/2020.02.18.953836

**Authors:** Sandra Goutte, Matías I. Muñoz, Michael J. Ryan, Wouter Halfwerk

## Abstract

In animal communication, receivers benefit from signals providing reliable information on signallers’ traits of interest. Individuals involved in conflicts, such as competition between rivals, should pay particular attention to cues that are ‘unfakeable’ by the senders due to the intrinsic properties of the production process. In bioacoustics, the best-known example of such ‘index signals’ is the relationship between a sender’s body size and the dominant frequency of their vocalizations. Dominant frequency may however not only depend on an animal’s morphology but also on the interaction between the sound production system and its immediate environment. Here, we experimentally altered the environment surrounding calling frogs and assessed its impact on the signal produced. More specifically, we altered water level, which forced frogs to float on the surface and tested how this manipulation affected the shuttling of air between the lungs and the vocal sac, and how this in turn impacted the calls’ dominant frequency. Our results show that frogs that are floating are able to fully inflate their lungs and vocal sacs, and that the associated change in airflow or air pressure is correlated with a decrease of call dominant frequency.

## 1. Introduction

Animal communication signals are generally presumed to convey information about the sender to the intended receiver(s). These signals must, on average, provide reliable cues regarding sender’s attributes, such as e.g. identity, size, motivation, or fecundity, in order to be maintained throughout evolution [1–3]. The evolutionary stability of these signalling systems is however challenged in the context of sexual selection, as mates and rivals often have conflicting interests in maximizing their fitness [4].

Senders may benefit from signals providing unreliable information about their traits, especially when these trait characteristics do not trigger the preferred response by the receivers. For example, across many taxa as well as sensory modalities, males have evolved signals that make them appear larger than they really are [5]. Receivers, on the other hand, may evolve counter-strategies when these so called “deceptive” signals impose a fitness cost to them. Those relying on signals that are “unfakeable” due to morphological or physiological constraints imposed on their production process (also known as ‘index signals’ [1,3,6]) ensure reliable (‘honest’) communication and avoid such cost. Characteristics of these signals thus provide reliable information on the signaller’s particular morphological or physiological traits [7].

The most widely used example of ‘honest’ communication, especially in the context of sexual selection, is the negative frequency-body size relationship that is often reported in bioacoustics [5–7]. Vocalizations’ frequency range is determined by the size of the sound producing organs, and individuals that differ in size tend to call or sing at different frequencies, providing receivers with a reliable cue about the size of their mate or opponent [6,8]. Not surprisingly, correlations between body size and frequency both within and between species abound [9]. The observation that bigger animals vocalize at lower frequencies compared to smaller animals can almost be viewed as a fundamental law of communication in vertebrates [2,6]. The relatively few, but clear exceptions to the rule are therefore likely to reflect strong historical sexual or natural selection pressure operating on acoustic frequency. In some taxa, signallers have evolved extreme morphologies, such as coiled trachea [6], or air-filled pouches to lower their vocalizations beyond the constraints imposed by their overall body size [5]. The slow evolutionary change of sound production structures’ morphology, however, is not the only channel animals may use to lower their vocalizations’ frequency.

Certain characteristics of display sites can directly influence signal production and transmission. Signallers often exploit the resonance properties of the structures from where they call [10,11]. For example, the sounds of signallers that call or chirp from burrows in the ground or hollow tree cavities are amplified [11,12]. The amplification is typically frequency-dependent, allowing signallers in some cases to lower the dominant frequency of their vocalizations. Call-site properties may also influence signal production more directly through the interaction with signallers’ morphology. A recent study on túngara frogs revealed, for example, that when the water level is experimentally lowered males are less able to inflate their lungs and therefore there is less air available to push from their lungs through their larynx into their vocal sac [13]. The consequence of this environmentally induced constraint is that males in shallower water called at lower amplitudes and with less complexity, which in turn influenced their attractiveness to females. Importantly, the impact of water level treatment appeared to be size-dependent [14]. Larger males were more influenced by water level treatment compared to smaller males. In fact, a binary choice experiment, playing calls of a large versus small male from opposite directions revealed that females preferred the calls of the larger male when these were recorded in deep water, and calls of the smaller males when recorded from shallow water [14].

Here, we re-examine the videos of the experiments described above [13] with the specific aim to link call site-induced signalling constraints to variation in dominant frequency. More specifically, we selected the trials in which males were fully floating, allowing for maximum signalling performance, and the trials in which the same males were clearly non-floating. We compared floating and non-floating trials to test whether males were constrained in vocal sac inflation when non-floating, and whether this was related to variation in dominant frequencies.

## 2. Material and Methods

The study was carried out on male túngara frogs, *Engystomops (= Physalaemus) pustulosus* collected in August 2014 from Soberanía National Park, near Gamboa, Republic of Panama. All individuals were released back to the site from which they were collected on the same night. All experiments with frogs were licensed and approved by STRI (IACUC permit: 2014-0805-2017) and the Autoridad Nacional del Ambiente de Panama (SE/A-82–14).

Male frogs were recorded individually in a hemi-anechoic chamber under IR-lighting. At the start of the experiment they were placed in a small cage consisting of a ring of evenly spaced nylon monofilament (diameter of 0.05 mm fishing line every 0.5 cm). The cage was placed in a pool (diameter of 50 cm) containing a tube that allowed the experimenter to either add (using a funnel) or subtract (using a 50 ml syringe) water, in order to manipulate water depth at the position of the frog. Males were stimulated with a low-amplitude chorus recording until they were readily calling for 1 min. We assessed male calling behaviour at four different water depths (0.25, 0.5, 1.0, and 2.0 cm). Each trial lasted for 1 min, followed by a 2-min break during which the water depth was altered (starting 1 min before the next trial).

We recorded male calling behaviour with a camera that was mounted on top of the cage (mini 1/4” CCTV camera; 2.8 mm lens; connected to a desktop PC). We recorded male calls with a microphone setup (G.R.A.S. 40 BF microphone amplified by 20 dB by G.R.A.S. 26 AC amplifier connected to an Avisoft 116Hm Ultrasound gate, G.R.A.S. Sound & Vibration A/S, Holte, Denmark) onto a desktop PC, using a sampling rate of 50 kHz. The microphone was placed at a 45° angle and at a distance of 50 cm from the frog. The microphone was calibrated prior to each experiment using a tone generator (G.R.A.S. 42 AB, 114 dB at 1 kHz).

We analysed the videos of male calling in our setup and scored whether males were completely floating or not. We selected for each trial three video stills from the beginning, middle and end of a call bout. We selected for each call the video still with the maximum inflation of the lungs as well as maximum inflation of the vocal sac. We used the programme ImageJ to measure from the width of the vocal sac at maximum inflation (in mm). Sound recordings were analysed in SASLab Pro (Avisoft Bioacoustics, Berlin, Germany). We selected three calls from the start, middle and end of a trial and measured the dominant frequency of each call.

We assessed whether floating affected vocal sac inflation and dominant frequency during calling in R (v.3.2.2). We constructed linear mixed models using the package *lme4* with a Gaussian distribution with identity link function. Significance of fixed effect (floating or non-floating) was assessed using likelihood (ML) ratio test. All models were tested for normality, overdispersion and heteroscedasticy. See [13] for a more detailed description of the setup, experimental procedures and analyses.

## 3. Results

We placed male túngara frogs in an experimental setup that allowed us to manipulate the water level while a focal individual was calling (see methods). Thus, males called either floating or with their legs or belly touching the ground. All twenty males we tested floated in the trials in which the water level was raised above 2 cm whereas none of these males were able to float when we lowered the water level to below 0.25 cm. During intermediate water level (0.5 and 1 cm) trials, males would often have their front or hindlegs touching the base of the setup. When the water level was raised we could clearly see males actively pumping air into their lungs until they had enough buoyancy to allow them to float freely on the water surface.

We compared trials in which males were fully floating with trials in which they were partially submerged (with either their front or hindlegs touching the base). Floating males were able to inflate their vocal sacs and thus their lungs to a greater extent during trials in which they could float (GLMM; n = 20 males; χ^2^ = 67.67; d.f. = 1; P < 0.001; Fig. 1). The increase in vocal sac inflation was accompanied by a decrease in dominant frequency (from 995 ± 307 Hz to 797 ± 63 Hz; Fig. 1.) as we found floating males’ calls had a lower dominant frequency compared to non-floating males (χ^2^ = 67.65; d.f. = 1; P < 0.001).

**Fig. 1.**
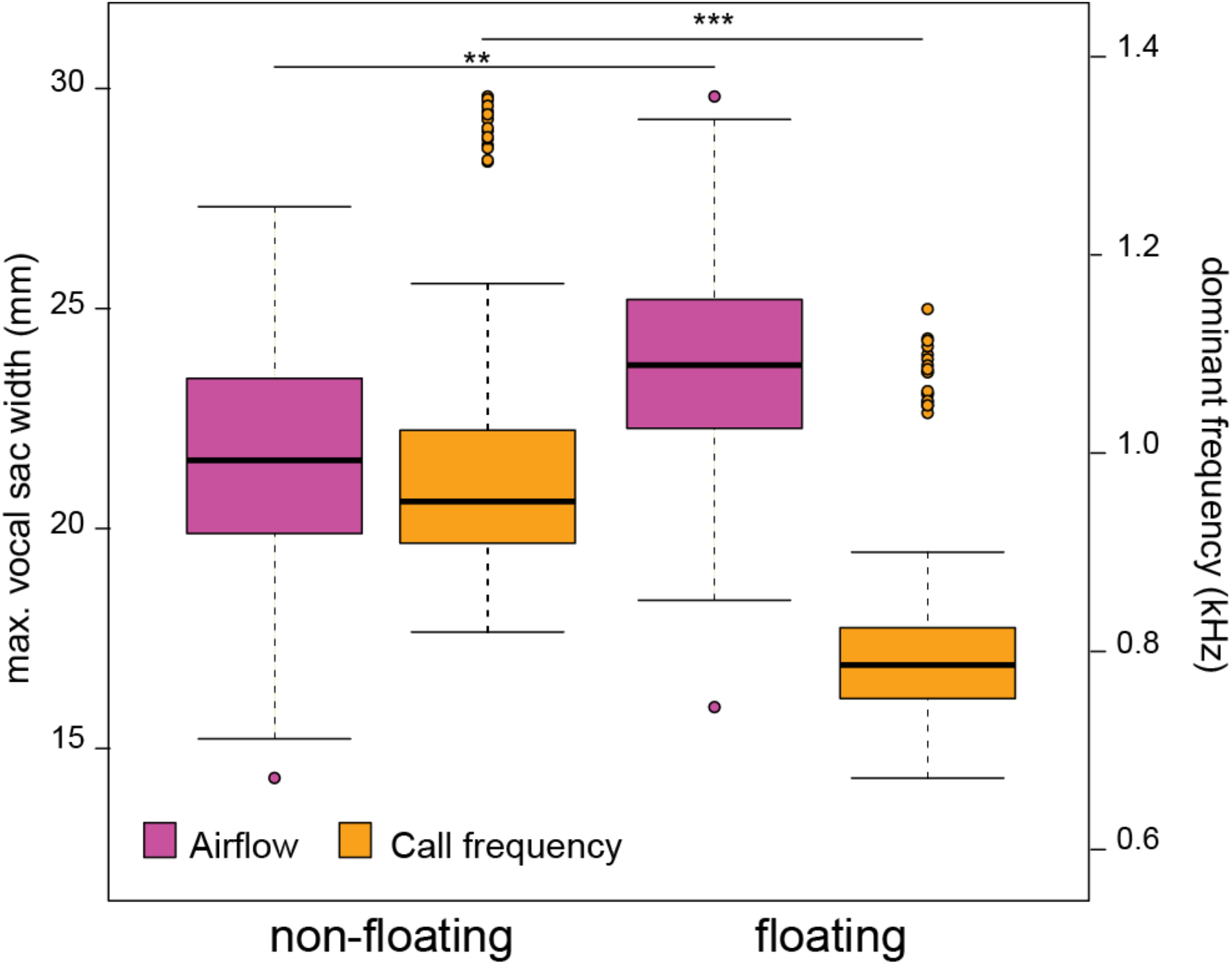
Effect of floating ability on vocal sac inflation and call dominant frequency

## 4. Discussion

The dominant frequency of vocalizations is generally assumed to be a reliable indicator of a signaller’s body size [15]. In this study, we show that the environment can directly influence signal production, thereby affecting dominant frequency, and hence alter signal reliability in reference to body size. We found that frogs vocalizing while floating on the water produced calls with lower frequencies compared to trials during which they did not float.

This environment-based modulation of call frequency may have important consequences on an individuals’ fitness. Multiple studies have shown that lower frequency calls are preferred by gravid females in frogs [16,17] in general, and specifically in túngara frogs [18,19,20,21]. Producing calls with lower frequencies thus confers a fitness benefit to the signaller by attracting more mates. In addition, in the species tested here, the lowering of the dominant frequency in floating males (about −200 Hz) approaches the 520 Hz peak sensitivity of the amphibian papilla (the organ responsible for hearing low-frequency airborne sounds) [22]. The decrease in frequency therefore might result in a greater detectability of floating males’ calls by conspecifics, which may facilitate recruiting females and repelling competitors over greater distances. Individual males can thus increase the reach and attractiveness of their calls by signalling from deeper puddles where they are able to float on the surface and inflate their lungs and vocal sacs to the fullest. In natural conditions, however, male túngara frogs are found vocalizing in both conditions (while floating and not floating), perhaps due to additional balancing selection pressures such as predation or reflecting the availability of ideal calling sites.

Our study demonstrating the effect of the external environment on the túngara frog’s dominant frequency has intriguing parallels to a study showing how this same frog’s internal environment influences call frequency. Kime et al. (2010) [23] showed that males treated with the hormone peptide arginine vasotocin (AVT) produce calls with higher frequencies, and that these calls are less attractive to females than the calls of control males. They speculated that AVT might influence motor output to cause more rapid airflow through the larynx and into the vocal sac.

Our results suggest that the properties of anuran call sites can have a strong influence on sexual selection. For example, in shallow water females may choose and rivals may respond stronger to smaller floating males over larger, non-floating ones as sound production is only constrained in larger males [14]. Display site properties may therefore drive signal evolution through biomechanical constraints on signal production.

